# Regulation of sexually dimorphic abdominal courtship behaviors in *Drosophila* by the *Tlx/tailless*-like nuclear receptor, *Dissatisfaction*

**DOI:** 10.1101/2021.08.09.455671

**Authors:** Julia C. Duckhorn, Jessica Cande, Mary C. Metkus, Hyeop Song, Sofia Altamirano, David L. Stern, Troy R. Shirangi

**Affiliations:** Villanova University, Department of Biology, 800 Lancaster Ave, Villanova, PA 19085; Janelia Research Campus, Howard Hughes Medical Institute, 19700 Helix Drive, Ashburn, VA 20147

## Abstract

Sexually dimorphic courtship behaviors in *Drosophila melanogaster* develop from the activity of the sexual differentiation genes, *doublesex (dsx)* and *fruitless (fru),* functioning with other regulatory factors that have received little attention. The *dissatisfaction* gene *(dsf)* encodes an orphan nuclear receptor homologous to vertebrate *Tlx* and *Drosophila tailless* that is critical for the development of several aspects of female- and male-specific sexual behaviors. Here, we report the pattern of *dsf* expression in the central nervous system and show that the activity of sexually dimorphic abdominal interneurons that co-express *dsf* and *dsx* is necessary and sufficient for vaginal plate opening in virgin females, ovipositor extrusion in mated females, and abdominal curling in males during courtship. We find that *dsf* activity results in different neuroanatomical outcomes in females and males, promoting and suppressing, respectively, female development and function of these neurons depending upon the sexual state of *dsx* expression. We posit that *dsf* and *dsx* interact to specify sex differences in the neural circuitry for dimorphic abdominal behaviors.

## Introduction

The neural circuitry for sex-specific behaviors in *Drosophila melanogaster* is currently understood to be specified during development by transcriptional regulators encoded by the *doublesex (dsx)* and *fruitless (fru)* genes [1]. The male isoforms of *dsx* and *fru* direct the differentiation of neurons in the adult male central nervous system that are required for performance of male courtship behaviors [2–6]. Conversely, the female-specific isoform of the *dsx* gene together with the absence of male-specific *fru* build the neural pathways that direct female-specific sexual behaviors [3,6–9].

Although *dsx* and *fru* are indispensable for sexual differentiation of the *Drosophila* nervous system, they act together with other regulatory genes that have been less well studied [10,11]. The *dissatisafaction* gene *(dsf),* for instance, which encodes an orphan nuclear receptor homologous to vertebrate *Tlx* and *Drosophila tailless,* is required for several aspects of female- and male-specific reproductive behaviors [12,13]. Females homozygous for null mutations of *dsf* rarely mate with courting males, and, if fertilized, are unable to lay eggs despite being able to ovulate [12]. Males mutant for *dsf* court females and males indiscriminately and are delayed in copulating with females at least in part due to deficits in abdominal bending during copulation attempts [12]. The *dsf* gene was first identified over twenty years ago, yet where and when *dsf* is precisely expressed in the *Drosophila* nervous system, which *dsf*-expressing neurons matter for courtship behavior, and how *dsf* contributes to the sexual differentiation of the nervous system are not known.

Here, we report *dsf*’s spatial and temporal expression in the central nervous system and discover a small group of sexually dimorphic *dsf* and *dsx* co-expressing interneurons in the abdominal ganglion called the DDAG neurons. Females and males have approximately eleven and three DDAG neurons per side, respectively, that exhibit sexually dimorphic projection patterns in the ventral nerve cord. Optogenetic activation and neural silencing experiments demonstrate that the DDAG neurons contribute to vaginal plate opening in virgin females, ovipositor extrusion in mated females, and abdominal curling in males during courtship. *Dsf* promotes the presence of female-specific DDAG neurons and the development of vaginal plate opening behavior in virgin females. In males, *dsf* acts in an opposite manner, suppressing the development of a single female-like DDAG neuron. Optogenetic activation of artificially induced female-like DDAG neurons in males drives an abdominal behavior in males that imitates vaginal plate opening or ovipositor extrusion in females. We show that *dsf* promotes or suppresses female-like development and function of the DDAG neurons depending upon the expression of the male isoform of *dsx.* Taken together, our results illustrate how the *dsf* gene regulates the development and function of sexually dimorphic neurons that mediate sexspecific abdominal behaviors.

## Results

### Expression of *dsf* in the *Drosophila* central nervous system

In a previous study [13], *dsf* expression in the *Drosophila* central nervous system (CNS) was detected by *in situ* hybridization in a small number of neurons located primarily in anterior regions of the pupal and adult protocerebrum. We reexamined *dsf* mRNA expression in the adult CNS by *in situ* hybridization chain reaction (HCR), a robust, quantitative, and sensitive method to detect mRNAs in whole-mounted tissues [14–17]. Using HCR probe-sets directed against *dsf*, we observed *dsf* mRNAs in the Canton S CNS in substantially more cells than originally reported (Figure 1A, B). In the brain, *dsf* expression was found in several broadly distributed clusters of cells located mostly on the anterior side, dorsal and ventral to the foramen of the esophagus. Hybridization signals were also detected in a few cells in the thoracic and abdominal neuromeres of the ventral nerve cord (VNC). The intensities and distributions of hybridization signals appeared similar between the sexes, and hybridization signals were not detected in flies null for the *dsf* gene (Figure 1C, D).

**Figure 1.**
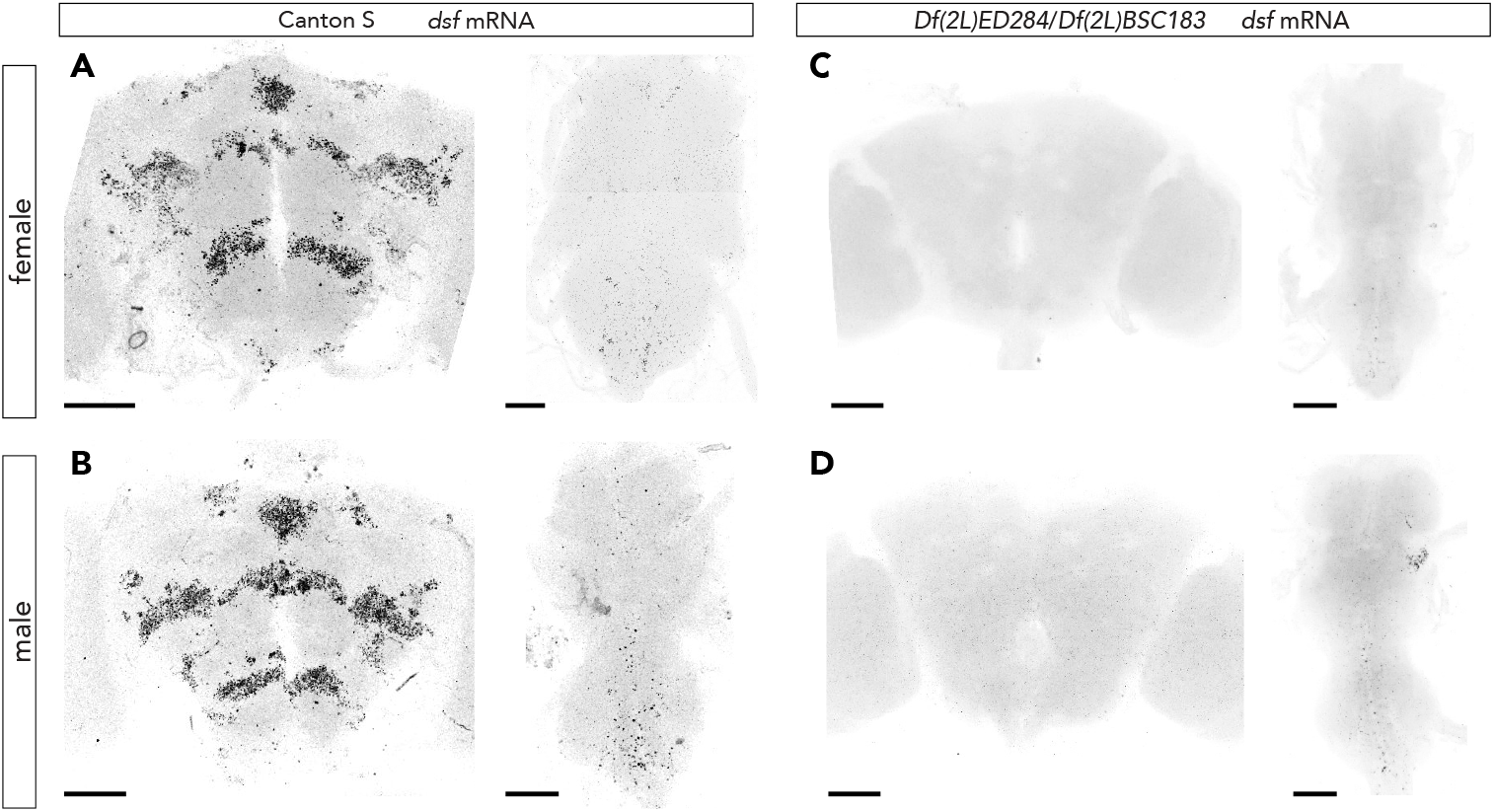
Expression of *dsf* mRNA in the adult *Drosophila* central nervous system. Confocal image of an adult Canton S female (A) and male (B) brain and VNC showing *dsf* mRNA expression labeled by *in situ* HCR using a probe-set directed against *dsf* transcripts. Hybridization signals are seen in several groups of cells in the brain and VNC. *dsf* mRNA expression was not detected in the CNSs of females (C) and males (D) carrying partially overlapping deficiencies that delete the *dsf* gene. Scale bar represents 50 μm.

To characterize the identity, anatomy, and function of *dsf*-expressing cells, we used CRISPR/Cas9-mediated homology-directed repair to insert the Gal4 transcriptional activator from yeast into the *dsf* translational start codon, generating a *dsf*^Ga14^ allele (Figure 2A). Insertion of Gal4 deleted all coding sequences within *dsf*’s first exon and the exon’s 5’ splice site, thereby creating a loss-of-function allele. We assessed whether *dsf*^Ga14^ reproduces the endogenous pattern of *dsf* transcripts by sequentially applying *in situ* HCR and immunohistochemistry to adult CNS tissues from flies heterozygous for *dsf*^Ga14^ and carrying *UAS-nls::gfp* to simultaneously detect *dsf* transcripts and GFP expression. We found an almost perfect overlap of the two markers in the adult brain and VNC of females and males (Figure 2B, C). All cells in the adult CNS labeled by *dsf*^Ga14^ co-expressed the pan-neuronal marker, ELAV, confirming that the *dsf*-expressing cells are neuronal (Supplemental Figure 1A–D).

**Figure 2.**
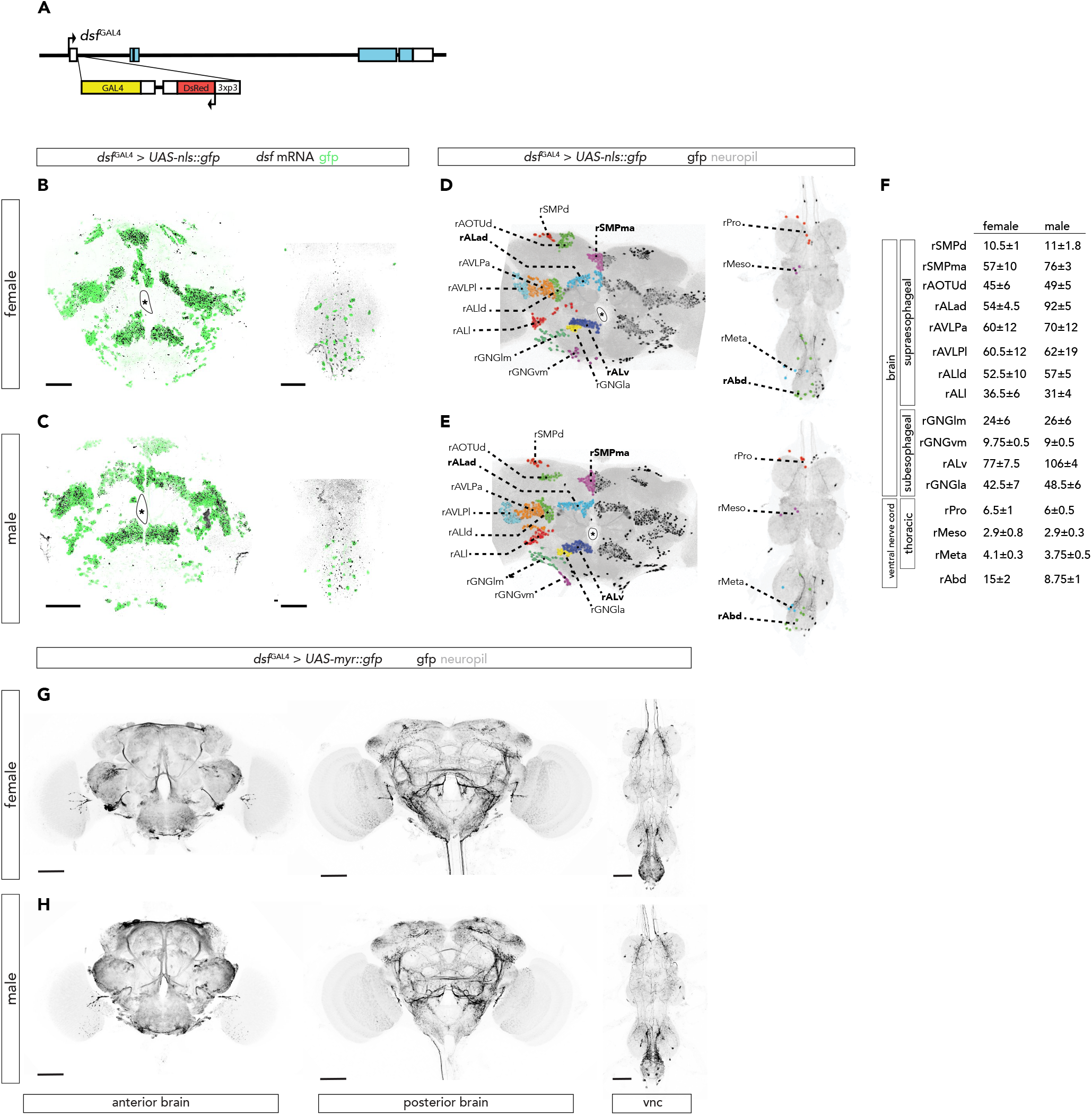
*dsf*^Ga14^ labels *dsf*-expressing neurons in the female and male adult central nervous system. (A) Design of the *dsf*^Ga14^ allele. Targeted insertion of the Gal4 coding sequence (yellow box) after *dsf*’s translational start codon in exon 1 allows expression of Gal4 wherever *dsf* is normally expressed. Each box represents an exon. Exonic regions colored in light blue and white represent *dsf* coding sequences and untranslated regions, respectively. (B, C) Confocal images of the adult brain, and third thoracic and abdominal ganglia of the VNC from *dsf*^Ga14^ > *UAS-nls::gfp* females (B) and males (C) showing co-localization of *dsf* mRNA (black) and GFP protein (green) labeled by combined *in situ* HCR and immunohistochemistry. *dsf* mRNA and GFP protein expression overlap in the brain. GFP-expressing cells in the VNC exhibit *in situ* HCR signals, however some hybridization signals in the VNC lack GFP expression. These hybridization signals are non-specific to *dsf*, as they persist in the VNCs of flies carrying deficiencies that delete the *dsf* locus (see Figure 1C, D). Scale bar represents 50 μm. (D, E) Confocal images of brains and VNCs from *dsf*^Ga14^ > *UAS-nls::gfp* females (D) and males (E) showing the distribution of GFP-expressing neurons in black and DNCad (neuropil) in light gray. Discrete groups of neurons were color-coded and categorized on one side of the brain according to standardized nomenclatures. Neuronal groups that exhibit sex differences in neuron number are labeled in bold. (F) Table showing the average number (±SD) of neurons in each *dsf*^Ga14^-expressing neuronal group (N=4 sides/group) in the brain and VNC of females and males. (G, H) Confocal images of brains and VNCs from *dsf*^Ga14^ > *UAS-myr::gfp* of females (G) and males (H) showing the projection and arborization patterns of *dsf*^Ga14^-expressing neurons in black and DNCad (neuropil) in light gray. Scale bar represents 50 μm.

We categorized *dsf*-expressing neurons as belonging to specific clusters of neurons and used standardized nomenclature [18,19] to assign names to each neuronal group based on their location in the cell body rind of the brain and VNC (Figure 2D–F). *dsf*-expressing neurons in the brain occur in twelve discernible groups on the left and right sides. Eight of the neuronal groups are in the supraesophageal zone of the brain and four are present in the subesophageal zone. All twelve groups of neurons are present in both sexes, but some groups *(i.e.,* rSMPma, rALad, and rALv) exhibit notable differences in neuron number between the sexes (Figure 2F). Additionally, females have a greater number of *dsf*-expressing neurons than males in the abdominal ganglion of the VNC (Figure 2F).

When driving a membrane-targeted GFP, *dsf*^Ga14^ labeled many different nerve bundles and synaptic neuropils present mostly on the posterior side of the adult brain and in the VNC (Figure 2G, H). No obvious sex differences in the projection patterns of *dsf*-expressing neurons were observed in the adult CNS. *dsf*^Ga14^ also labeled subsets of neurons in the late 3^rd^ instar larval and pupal (Supplemental Figure 1E–J) CNS.

*Dsf* mutant females and males were reported to display abnormalities in motoneuron innervation on muscles of the uterus and ventral abdomen, respectively [12]. We did not detect *dsf*^Ga14^ activity in motoneurons (or peripheral sensory neurons) at any stage that we examined, nor in muscles of the ventral abdomen or uterus. To further confirm that *dsf*^Ga14^ accurately labeled all *dsf*-expressing neurons in the abdominal ganglion, we sequentially detected *dsf* mRNAs and GFP and ELAV protein expression in CNS tissue from *dsf*^Ga14^ > *myr::gfp* flies. *Dsf* hybridization signals were detected exclusively in ELAV-expressing neurons that coexpressed GFP, indicating that all *dsf*-expressing neurons are targeted by the *dsf*^Ga14^ allele (Supplemental Figure 1K). These data suggest that the neuromuscular junction phenotypes of *dsf* mutant flies [12] may result from cell-non-autonomous actions of *dsf*. Taken together, we conclude that *dsf* is expressed in several subsets of neurons broadly distributed across the female and male CNS of adult, pupal, and larval stages, and that Gal4 expression from the *dsf*^Ga14^ allele faithfully recapitulates the *dsf* wild-type expression pattern.

### A sexually dimorphic group of *dsf* and *dsx* co-expressing neurons contribute to female- and male-specific abdominal courtship behaviors

*dsf* contributes to courtship behaviors [12] that are also influenced by *dsx* gene function or *dsx*-expressing neurons [6–8,20]. We reasoned that *dsf*-expressing neurons relevant for courtship behaviors may co-express *dsx*. We therefore genetically intersected *dsf*^Ga14^ with *dsx*^LexA^ [20], which expresses *LexA::p65* in all *dsx*-expressing cells. *Dsx^LexA^* was used to drive a *LexAop-* controlled Flp recombinase, which excised a transcriptional stop cassette from an upstream activating sequence (UAS)-myr::GFP transgene driven by *dsf*^Ga14^. This intersection (*dsf*^Ga14^ ⋂ *dsx*^LexA^) labeled a small group of sexually dimorphic interneurons (hereafter referred to as the *<u>d</u>sf-<u>d</u>sx* <u>a</u>bdominal <u>g</u>anglion, or DDAG, neurons) in the abdominal ganglion of both sexes (Figure 3A, B). Males have three DDAG neurons on the left and right sides that arborize locally within the abdominal neuropil and whose cell bodies are located on the ventral side of the VNC. Females possess approximately eleven DDAG interneurons, many of which are located on the dorsal side of the VNC. It is currently unclear if females have homologs of the DDAG neurons found in males, or if females and males have entirely sex-specific DDAG neurons. The DDAG neurons of females arborize extensively within the abdominal ganglion, and some neurons extend neurites anteriorly to innervate all three thoracic neuropils. The intersection between *dsf*^Ga14^ and *dsx^LexA^* does not appear to label any other cells in the brain or VNC (Figure 3A, B), nor any non-neuronal somatic tissues in the abdomen or elsewhere in the adult (data not shown).

**Figure 3.**
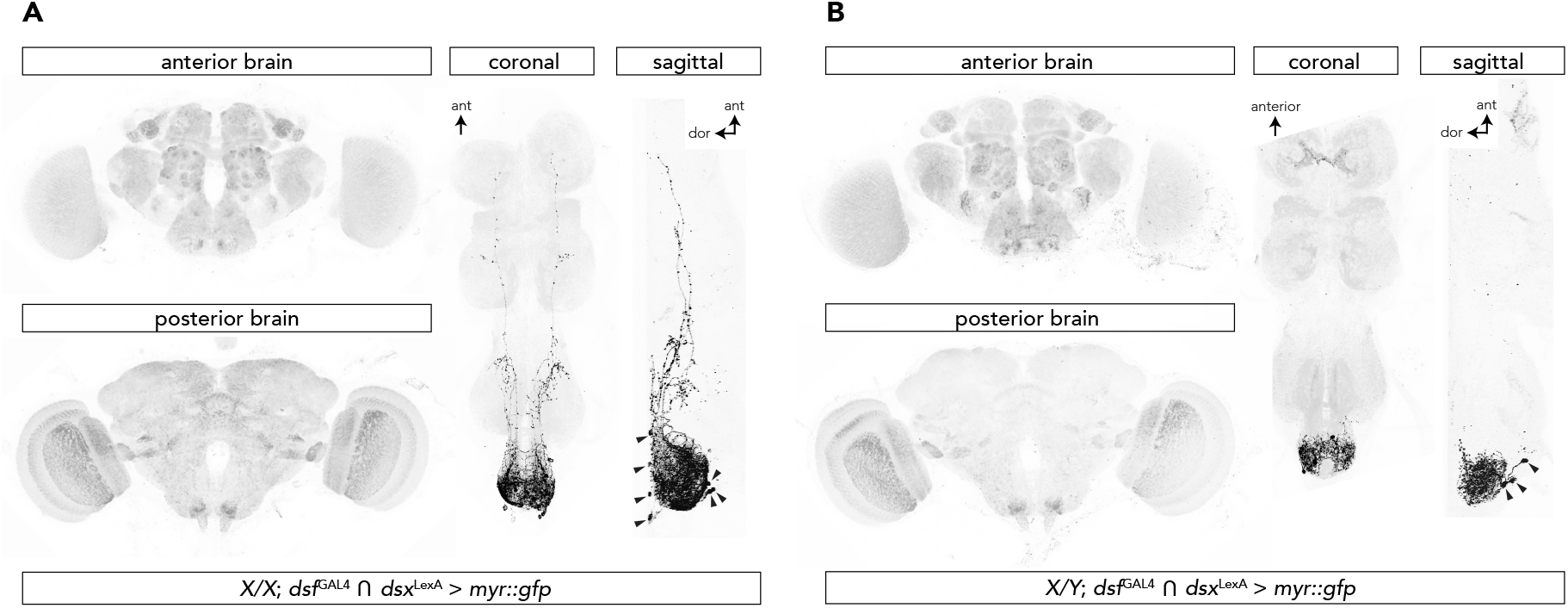
A sexually dimorphic group of *dsf* and *dsx* co-expressing interneurons in abdominal ganglion. The intersection of *dsf*^Ga14^ and *dsx*^LexA^ targets the sexually dimorphic DDAG neurons in the abdominal ganglion of females (A) and males (B). GFP-expressing neurons and DNCad (neuropil) are shown in black and light gray, respectively. *dsf*^Ga14^ ⋂ *dsx*^LexA^ labels no other neurons in the VNC or brain. Arrowheads point to the cell bodies of the DDAG neurons. Not all cell bodies are visible.

To determine whether the DDAG neurons contribute to courtship behaviors, the *dsf^Ga14^* ⋂ *dsx*^LexA^ intersection was used to target the expression of the red light-gated cation channel *CsChrimson* [21] to activate the DDAG neurons. When stimulated with red light, *dsf*^Ga14^ ⋂ *dsx^LexA^* > *CsChrimson* virgin females adjusted the posture and length of their abdomen and opened their vaginal plates, partially exposing their ovipositor (Supplemental Video 1; Figure 4A). Virgin females normally open their vaginal plates if they are receptive to courting males [22,23] in contrast to mated females, which signal their unwillingness to mate by extruding their ovipositor [22,24,25]. The length of the female’s abdomen increases during both behaviors, but by a greater amount during an ovipositor extrusion (Figure 4B). To confirm that photoactivation of the DDAG neurons in virgin females triggered an opening of the vaginal plates and not an ovipositor extrusion, we measured the change in abdomen length before and during photoactivation and compared it to the change in abdomen length when virgin females open their vaginal plates or when mated females extrude their ovipositor. Optogenetic activation of the DDAG neurons in virgin females induced a change in abdomen length more similar to that measured when virgin females open their vaginal plates during courtship than when mated females extrude their ovipositor (Figure 4B). Virgin *dsf^Ga14^* ⋂ *dsx*^LexA^ > *CsChrimson* females opened their vaginal plates immediately and only during the photoactivation period (Figure 4C), and quantitatively similar behaviors were observed across a range of stimulus intensities (Figure 4D, E). In mated females, however, photoactivation of the DDAG neurons induced ovipositor extrusion (Supplemental Video 2; Figure 4A, B), suggesting that female mating status alters the function of the DDAG neurons. Female receptivity in *Drosophila* is modified after mating by Sex Peptide (SP), a male seminal protein that is transferred to the female reproductive system during copulation [26]. When mated with SP null males, *dsf^Ga14^* ⋂ *dsx^LexA^* > *CsChrimson* females opened their vaginal plates instead of extruding their ovipositor during photoactivation (Supplemental Video 3; Figure 4B). These data suggest that the DDAG neurons contribute to virgin or mated female abdominal behaviors depending on the mating status of the female.

**Figure 4.**
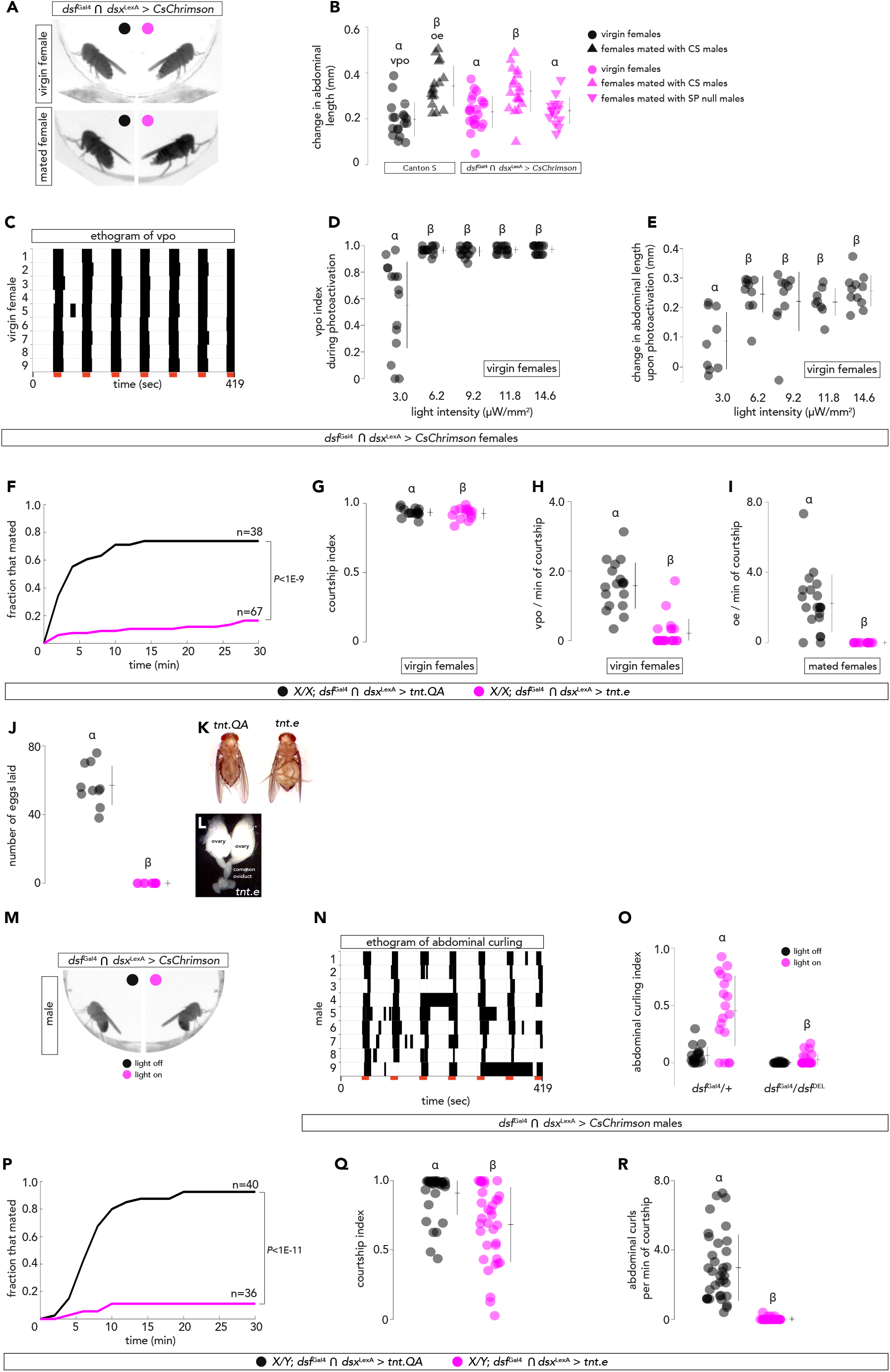
Activity of the DDAG neurons is sufficient and necessary for female- and male-specific abdominal courtship behaviors. (A) A still frame image of a *dsf*^Ga14^ ⋂ *dsx*^LexA^ > *CsChrimson* virgin and mated female, immobilized from decapitation, before (left) and during (right) illumination with 14.6 μW/mm^2^ of red light to photoactivate the DDAG neurons. The virgin female opens her vaginal plates, whereas the mated female extrudes her ovipositor. (B) The change in abdominal length upon an opening of the vaginal plates in virgin females and an ovipositor extrusion in mated females was measured and compared to the change in abdominal length induced upon photoactivation of the DDAG neurons in virgin females and females that mated with Canton S or Sex Peptide (SP) null males. vpo, vaginal plate opening; oe, ovipositor extrusion. (C) An ethogram of vaginal plate opening behavior of nine *dsf*^Ga14^ ⋂ *dsx*^LexA^ > *CsChrimson* virgin females during seven 15-sec bouts of photoactivation (red bars). Black bars indicate when the female opens her vaginal plates. (D, E) Fraction of time a virgin female opens her vaginal plates during a 15-s bout of photoactivation (*i.e*., vpo index) and the change in abdominal length upon photoactivation (E) are largely unchanged across a variety of light intensities. (F–H) Virgin females with inhibited DDAG neurons using tetanus neurotoxin (magenta) are unreceptive to active male courtship and exhibit a reduced frequency of vaginal plate opening behaviors relative to the control (black). (F) Fraction of virgin females that mated with a Canton S male over 30 minutes. (G) Fraction of time a Canton S male spent courting a virgin female during 5 minutes of observation (*i.e.*, courtship index). (H) Number of times a virgin female opened her vaginal plates per minute of active courtship by a Canton S male. (I) Number of times a mated female extruded her ovipositor per minute of active courtship by a Canton S male. (J) Number of eggs laid by a female 20-22 hours after mating. (K) Mated females with inhibited DDAG neurons *(tnt.e)* appear gravid due to egg retention relative control *(tnt.QA)* females. (L) Eggs are observed in the oviducts of a mated female with inhibited DDAG neurons, indicating that they ovulate. (M) A still frame image of a decapitated *dsf*^Ga14^ ⋂ *dsx*^LexA^ > *CsChrimson* male before (left) and during (right) illumination with 14.6 μW/mm^2^ of red light to photoactivate the DDAG neurons. Photoactivation results in abdominal curling without extrusion of the male’s terminalia. (N) An ethogram of abdominal curling of nine *dsf*^Ga14^ ⋂ *dsx*^LexA^ > *CsChrimson* males during seven 15-sec bouts of photoactivation (red bars). Black bars indicate when the male curls his abdomen. (O) The fraction of time a *dsf*^Ga14^ ⋂ *dsx*^LexA^ > *CsChrimson* male curls his abdomen (*i.e*., abdominal curling index) during bouts of photoactivation (magenta) and darkness (black) in a *dsf* heterozygous (*dsf*^Ga14^/+) and *dsf* mutant (*dsf*^Ga14^/*dsf*^Del^) background. (P-R) Males with inhibited DDAG neurons using tetanus neurotoxin (magenta) exhibit reduced mating rates and frequency of abdominal curls per minute of active courtship relative to the control (black). (D) Fraction of males that mated with a Canton S virgin female over 30 minutes. (E) Courtship index of males courting Canton S virgin females. (F) Number of abdominal curls per minute of active courtship. (B, D, E, G–J, O, Q, R) show individual points, mean, and SD. Significance (P<0.05) was measured in (B, D, O) using a one-way ANOVA with a Tukey-Kramer test for multiple comparisons, in (G–J, Q, R) using a Rank Sum test, and in (F, P) using a Logrank test. Same letter denotes no significant difference.

We next tested whether the activity of the DDAG neurons is required for vaginal plate opening in virgin females and ovipositor extrusion in mated females. We used *dsf^Ga14^* ⋂ *dsx*^LexA^ to drive the expression of tetanus neurotoxin light chain *(tnt.e)* [27] to suppress the function of the DDAG neurons in virgin females. Compared to a control that expressed an inactive form of the neurotoxin *(tnt.QA*), virgin females that expressed *tnt.e* in the DDAG neurons mated infrequently with courting males (Figure 4F) despite being vigorously courted (Figure 4G), and exhibited a marked reduction in the frequency of vaginal plate opening during courtship (Figure 4H). Virgin females with inhibited DDAG neurons did not display behaviors indicative of mated females, such as ovipositor extrusion. Control females and females with inhibited DDAG neurons that eventually mated were tested 24 hours later in courtship assays with wildtype males. Mated *dsf*^Ga14^ ⋂ *dsx*^LexA^ > *tnt.e* females extruded their ovipositor to courting males less frequently compared to mated *tnt.QA*-expressing control females (Figure 4I). Additionally, mated females with inhibited DDAG neurons produced and released eggs into the oviduct (Figure 4J) but failed to lay any eggs (Figure 4K) and eventually appeared gravid (Figure 4L). *Dsf* null females lack synapses on the uterine wall (Supplementary Figure 2A–C and [12]), which correlates with their inability to lay eggs. However, inhibition of the DDAG neurons with *tnt.e* did not affect the presence or gross morphology of uterine synapses compared to *tnt.QA-* expressing control females (Supplemental Figure 2D, E). A subset of DDAG neurons may thus contribute to egg laying by innervating uterine motoneurons. Together, these results demonstrate that activity of the DDAG neurons is required for vaginal plate opening in virgin females, and ovipositor extrusion and later steps of egg laying in mated females.

In males, we observed that photoactivation of the DDAG neurons induced abdomen bending (Supplemental Video 4; Figure 4M–O). During *Drosophila* courtship, males attempt to copulate with females by curling their abdomen ventrally by ~180° to achieve contact between genitalia. Males with inhibited DDAG neurons (*dsf*^Ga14^ ⋂ *dsx*^LexA^ > *tnt.e)* actively courted Canton S virgin females, but were strongly reduced in successful mating rates relative to control (*dsf*^Ga14^ ⋂ *dsx*^LexA^ > *tnt.QA)* males (Figure 4P, Q). Additionally, in comparison with controls, males with inhibited DDAG neurons showed markedly reduced frequency of abdomen bending per minute of active courtship (Figure 4R). We conclude that the DDAG neurons of females and males influence sex-specific abdominal behaviors during courtship.

### Sexually dimorphic anatomy and function of the DDAG neurons results from *dsxM*-mediated defeminization

We next sought to determine how the dimorphic development of the DDAG neurons is genetically regulated. *Dsf*^Ga14^ ⋂ *dsx*^LexA^ > *myr::gfp* was used to visualize the DDAG neurons as before, but now we used a validated UAS-regulated short hairpin/miRNA (ShmiR) targeting *dsx* [28,29] to deplete male-specific *dsx (dsxM)* or female-specific *dsx (dsxF)* transcripts in *dsf*-expressing cells of both sexes. Depletion of male-specific *dsx* (*dsx*M) transcripts in *dsf*-expressing cells resulted in a near complete anatomical feminization of the DDAG neurons in males (compare Figure 5B to 5A); males gained approximately eight DDAG interneurons (Figure 5M) with an arborization pattern similar to that of the DDAG neurons of females (compare Figure 5B to 5G). Knock-down of female-specific *dsx (dsxF)* transcripts, however, did not result in any obvious changes in DDAG neuron morphology or number (compare Figure 5H to 5G; Figure 5N). These results indicate that sex differences in the DDAG neurons result at least in part from *dsx*M-mediated defeminization, whereby *dsx*M removes a set of neurons normally found in females. In the absence of sexual differentiation *(i.e.,* without *dsx* function), the DDAG neurons develop the same in both sexes but in the likeness of DDAG neurons normally observed in females.

**Figure 5.**
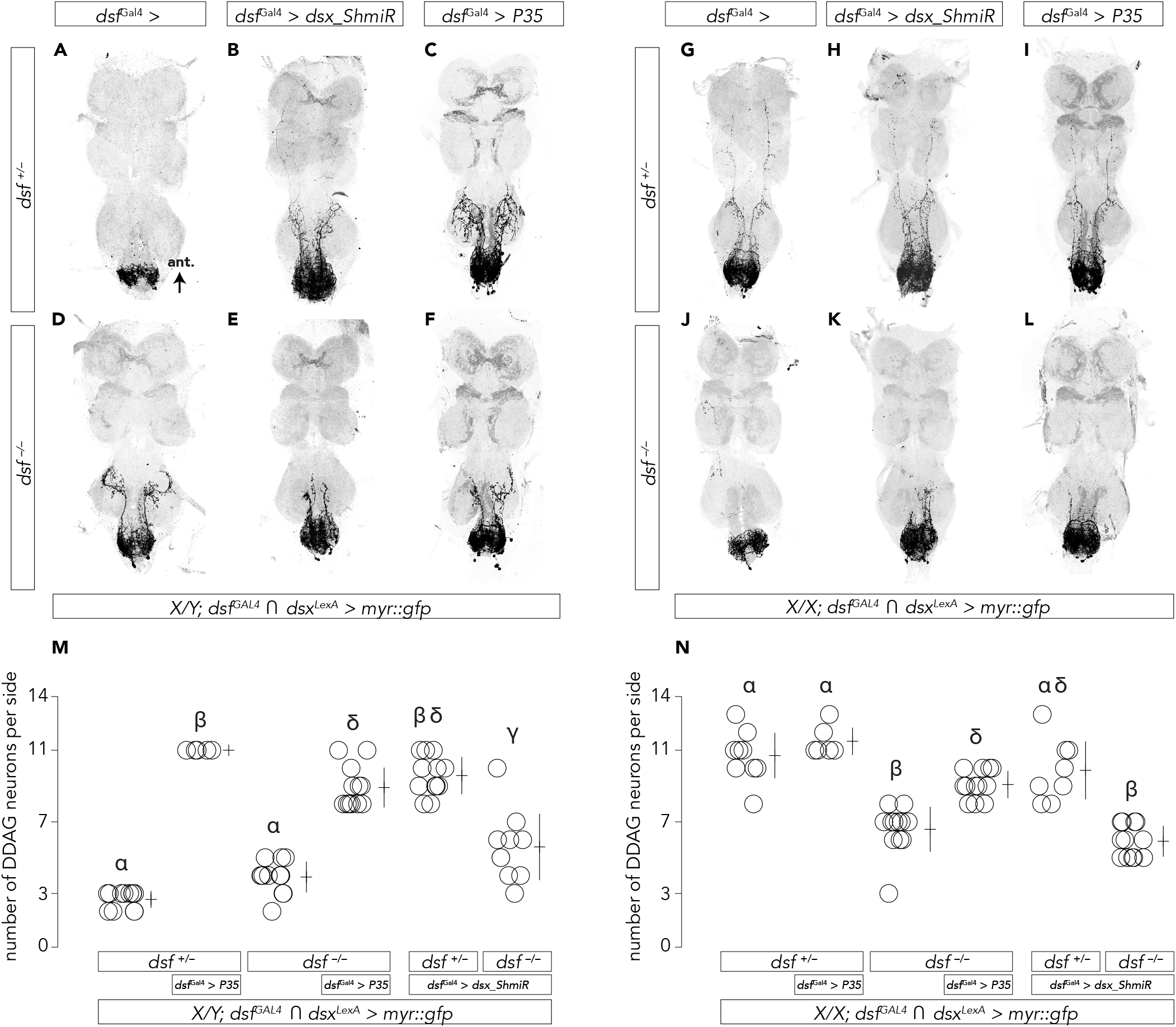
*dsx* and *dsf* influence sexually dimorphic DDAG neuron number through regulation of cell death. Exemplary confocal images of male (A-F) and female (G-L) VNCs from the genotypes analyzed in (M) and (N), respectively. The DDAG neurons were visualized in *dsf*^Ga14^ ⋂ *dsx*^LexA^ > *myr::gfp* flies. GFP-expressing neurons and DNCad (neuropil) are shown in black and light gray, respectively. (M, N) The number of DDAG neurons per side in VNCs of males (M) and females (N). (M, N) show individual points, mean, and SD. Significance (P<0.05) was measured in using a one-way ANOVA with a Tukey-Kramer test for multiple comparisons. Same letter denotes no significant difference.

To test if *dsxM* removes female-specific DDAG neurons by apoptosis, we used *dsf^Ga14^* to ectopically express the baculoviral caspase inhibitor, *P35* [30], in the DDAG neurons. Inhibition of cell death in males resulted in the gain of approximately eight DDAG neurons (Figure 5M) that exhibited grossly female-like projection patterns (compare Figure 5C to 5G). The morphology of the resurrected DDAG neurons in males is not fully feminized, however, most likely due to expression of *dsx*M. Inhibition of cell death did not change the number of DDAG neurons in females (Figure 5N; compare Figure 5I to 5G). These data demonstrate that *dsx*M is required for apoptosis of a subset of DDAG neurons in males that otherwise survive in females due to the absence of *dsx*M activity.

To determine whether the depletion of *dsx* activity influences the function of the DDAG neurons, we photoactivated the DDAG neurons in females and males in which *dsx_ShmiR* was driven by *dsf*^Ga14^. Photoactivation triggered the opening of the vaginal plates in females, similar to control females that lacked the *UAS-dsx_ShmiR* transgene (Supplemental Video 5; Figure 6A), consistent with the absence of an obvious neuroanatomical phenotype in the DDAG neurons of *dsf*^Ga14^ > *UAS-dsx_ShmiR* females. In contrast, photoactivation failed to induce abdomen curling in males, but instead triggered a behavior in which the males extended their abdomen and extruded their terminalia (*i.e.*, the terminal appendage consisting of genital and anal structures) during the illumination bout (Supplemental Video 6; Figure 6B, C). Terminalia extrusion was never observed when the DDAG neurons were photoactivated in control males (Supplemental Video 4; Figure 6C), suggesting that this behavior is gained in males with feminized DDAG neurons. Although courting males normally extrude their terminalia as they curl their abdomen during a copulation attempt, the terminalia extrusions resulting from photoactivation of feminized DDAG neurons were accompanied with postural changes in the abdomen that phenocopy female-like abdominal courtship behaviors like vaginal plate opening or ovipositor extrusion (Supplemental Video 6; Figure 6B).

**Figure 6.**
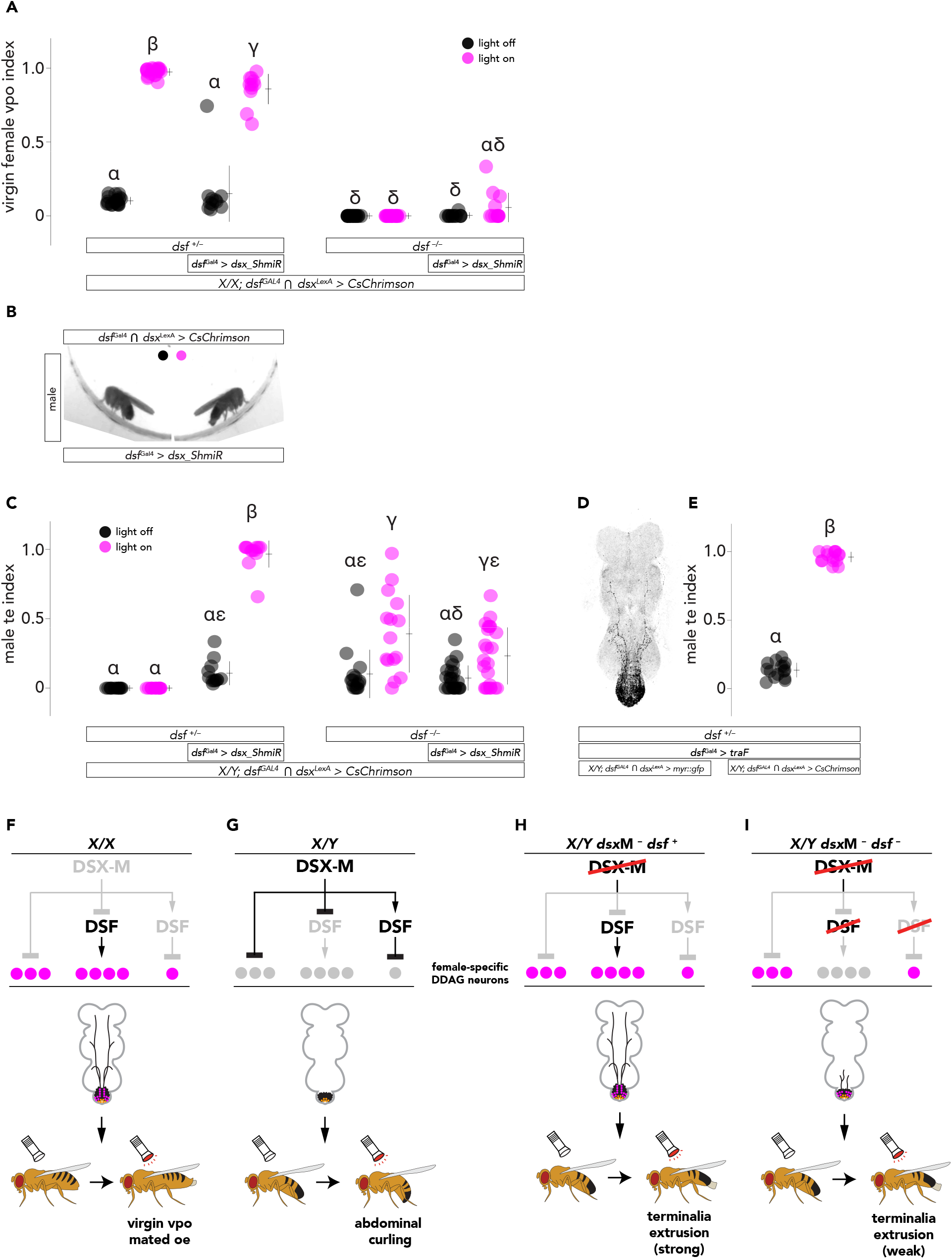
*dsf* promotes and suppresses female-type function of the DDAG neurons depending upon the expression of *dsx*. (A) The fraction of time a virgin female expressing *CsChrimson* in the DDAG neurons spends with her vaginal plates open (*i.e.*, vpo index) during darkness (black) and during 15-s bouts of photoactivation (magenta) with 14.6 μW/mm^2^ of red light. (B) A still frame image of a decapitated *dsf*^Ga14^ > *UAS-dsx_ShmiR* male before (left) and during (right) photoactivation of the DDAG neurons with 14.6 μW/mm^2^ of red light performing a terminalia extrusion. (C) The fraction of time a male expressing *CsChrimson* in the DDAG neurons spends extruding his terminalia (*i.e.*, terminalia extrusion, or te, index) during darkness (black) and during 15-s bouts of photoactivation (magenta) with 14.6 μW/mm^2^ of red light. (D) An exemplary VNC from a *tra*F-expressing male exhibiting fully feminized DDAG neurons. (E) The te index of a *tra*F-expressing male before (black) and during (magenta) photoactivation of the DDAG neurons with 14.6 μW/mm^2^ of red light. (F, G) Model for *dsx* and *dsf* function in regulating the number of female-specific DDAG neurons in wild-type females (F) and males (G) and the resulting neuroanatomical and behavioral outcomes. Female-specific DDAG neurons are shown in magenta. This model assumes that females share the DDAG neurons found in males, which are shown in orange. (H, I) Loss of *dsf* activity in males with depleted *dsx*M expression is predicted to result in a loss of four female-specific DDAG neurons and reduced levels of terminalia extrusion upon photoactivation of the DDAG neurons. (A, C, E) show individual points, mean, and SD. Significance (P<0.05) was measured using a one-way ANOVA with a Tukey-Kramer post hoc test for multiple comparisons. Same letter denotes no significant difference.

To confirm that these artificially induced terminalia extrusions result from a feminization of the DDAG neurons, we constructed *dsf^Cal4^* ⋂ *dsx^LexA^* > *CsChrimson* males that included a UAS-regulated female-specific *transformer (traF)* transgene [31], as an alternative strategy to feminize the DDAG neurons (Figure 6D). When stimulated with red light, the DDAG neurons of *dsf^Ga14^* > *UAS-traF* males triggered extrusions of the terminalia in ways and at levels similar to what was observed for *dsf^Cal4^* > *UAS-dsx_ShmiR* males (Figure 6E).

We next sought to determine how depletion of *dsx* transcripts in the DDAG neurons may affect the courtship behaviors of females and males. Using *dsf*^Ga14^ to drive the expression of *dsx_ShmiR*, we found that reduction of *dsx*F in the DDAG neurons of females did not affect mating rates with wild-type males (Supplemental Figure 3A), nor the frequency of vaginal plate openings per minute of courtship (Supplemental Figure 3B), but *dsf*^Ga14^ > *dsx_ShmiR* females laid fewer eggs (Supplemental Figure 3C) than control females. Depletion of *dsx*M transcripts in the DDAG neurons of males led to reduced mating rates with wild-type virgin females (Supplemental Figure 3D) and reduced frequency of abdominal bends during courtship (Supplemental Figure 3E). *Dsf*^Ga14^ > *dsx_ShmiR* males were never observed to extrude their terminalia during courtship. Taken together, these results indicate that male-specific *dsx* defeminizes the anatomy and function of the DDAG neurons in part by directing the cell death of female-specific DDAG neurons that contribute to female-specific abdominal courtship behaviors.

### *dsf* exerts opposite effects on the development and function of the DDAG neurons in females and males depending upon the expression of *dsxM*

Females carrying loss-of-function mutations in *dsf* mate infrequently with courting males, and *dsf* mutant males exhibit reduced copulation rates that correlate with abnormalities in abdomen bending during courtship [12]. Our results demonstrate that the DDAG neurons influence most courtship traits altered in *dsf* mutants. To determine whether *dsf* gene function is required specifically in the DDAG neurons for wild-type abdominal courtship behaviors, we depleted *dsf* expression in the DDAG neurons using *dsx*^Ga14(Δ2)^ [32] and a validated UAS-controlled *dsf_ShmiR.* Compared to controls that carried only either the Gal4 or UAS transgenes, *dsx*^Ga14(Δ2)^ > *UAS-dsf_ShmiR* virgin females mated infrequently with courting Canton S males, displayed a reduction in the frequency of vaginal plate opening during active male courtship, and retained their eggs (Supplemental Figure 3F–H). Depletion of *dsf* transcripts in the DDAG neurons of males resulted in reduced mating rates with Canton S virgin females and a reduced frequency of abdomen bends during courtship (Supplemental Figure 3I, J). These data suggest that *dsf* function is required in the DDAG neurons for female- and male-specific abdominal courtship behaviors that are altered in *dsf* mutants.

We next asked whether *dsf* influences the development of the DDAG neurons. We used *dsf*^Ga14^ ⋂ *dsx*^LexA^ > *myr::gfp* to visualize and compare the DDAG neurons of a *dsf* loss-of-function mutant (*dsf*^Ga14^/*dsf*^De1^) with a heterozygous control (*dsf*^Ga14^/+). Control females have approximately eleven bilaterally paired DDAG neurons on the left and right sides of the VNC, about four of which are lost in the absence of *dsf* function (Figure 5N). Additionally, the neurites of the DDAG neurons that normally project to the thoracic ganglia in control females were substantially reduced in *dsf* mutant females (compare Figure 5J to 5G). *Dsf* may influence DDAG neuron number by promoting the survival of a subset of DDAG neurons in females that would otherwise undergo apoptosis in the absence of *dsf* activity. Indeed, the number of DDAG neurons in *dsf* mutant females was partially restored when *dsf*^Ga14^ was used to drive the expression of baculoviral *P35* (Figure 5N; compare Figure 5L to 5J). Similar results were observed in *dsf* mutant males (Figure 5M; compare Figure 5F to 5D).

In males, the loss of *dsf* resulted in a gain of a single bilateral pair of DDAG neurons in the dorsal abdominal ganglion (Figure 5M; compare Figure 5D to 5A). This neuron extends a neurite shaped like a handlebar moustache anteriorly to innervate the third thoracic neuropil (Figure 5D). Both the neuron’s location on the dorsal side and its gross projection pattern are similar to DDAG neurons found in females, suggesting that the loss of *dsf* activity in males may have partially feminized the DDAG neurons. To determine how the loss of *dsf* activity may affect the function of the DDAG neurons, we photoactivated the DDAG neurons in *dsf* mutant females and males *(dsf^Ga14^/dsf^Del^* ⋂ *dsx*^LexA^ > *CsChrimson*). In contrast to control (*dsf*^Ga14^ ⋂ *dsx*^LexA^ > *CsChrimson)* females, photoactivation of the DDAG neurons in *dsf* mutants did not elicit vaginal plate opening (Supplemental Video 7; Figure 6A) nor abdominal curling in males (Figure 5O), but *dsf* mutant males extruded their terminalia during bouts of illumination (Supplemental Video 8; Figure 6C). The fraction of time *dsf* mutant males extruded their terminalia during an illumination bout was variable, and some *dsf* mutant males extruded their terminalia even after the illumination period had ended (Figure 6C). These results demonstrate that *dsf* promotes the presence of about four female-specific DDAG neurons in females, but functions in an opposite manner in males, suppressing the presence of a single female-like DDAG neuron whose activity is associated with a feminized abdominal behavior, *i.e.*, terminalia extrusion.

These observations support a model where *dsf* regulates the development of the DDAG neurons depending upon expression of *dsx*M. In females, when *dsx*M is absent, *dsf* promotes the survival of four female-specific DDAG neurons that likely contribute to female-specific abdominal courtship behaviors (Figure 6F). In males, *dsx*M blocks *dsf* function in promoting the presence of these female-specific neurons (Figure 6G). *Dsx*M defeminizes the DDAG neurons also by directly removing three female-specific neurons and by activating *dsf*’s role in suppressing the presence of a single female-like DDAG neuron (Figure 6G). In this model, loss of *dsf* function in males with reduced *dsx*M expression is predicted to result in the loss of four female-like DDAG neurons and reduced levels of photoactivated terminalia extrusion behavior relative to males with only depleted *dsx*M transcripts (Figure 6H, I).

Indeed, removal of *dsf* function in males with reduced *dsx*M expression led to a loss of about four DDAG neurons and a reduction of anteriorly-projecting neurites compared to control flies with only reduced *dsx*M expression (Figure 5M; compare Figure 5B to 5E). Removal of *dsf* activity in females with or without *dsx*F activity caused a similar phenotype (Figure 5N; compare Figure 5G to 5J, and Figure 5H to 5K). Additionally, males with reduced *dsx*M expression extruded their terminalia upon DDAG photoactivation at lower levels when *dsf* function was removed (Figure 6C). Photoactivation of the DDAG neurons in *dsf* mutant females did not induce vaginal plate opening behavior with or without depletion of *dsx*F expression (Figure 6A). These results suggest that *dsf* acts as both a ‘pro-female’ and ‘anti-female’ factor for DDAG development and function depending upon the expression of *dsx*M.

## Discussion

We have discovered a sexually dimorphic group of *dsf* and *dsx* co-expressing abdominal interneurons called the DDAG neurons that contribute to vaginal plate opening in virgin females, ovipositor extrusion in mated females, and abdominal curling in males during courtship. We provide evidence that male-specific *dsx* directs the dimorphic development of the DDAG neurons in part through regulation of *dsf* activity. Depending upon the absence or presence of *dsx*M, *dsf* promotes the development of female-type DDAG neurons that regulate the opening of the vaginal plates in females, but acts in an opposite manner in males, suppressing the development of female-like DDAG neurons and abdominal behaviors. Several groups of *dsf*-expressing, *dsx*-non-expressing neurons in the brain also exhibit sex differences in cell number. These neurons likely co-express *fru* and may contribute to other sex-specific behaviors altered in *dsf* mutants.

Neural circuits for vaginal plate opening and ovipositor extrusion and their regulation by the female’s mating status have been elucidated recently [22–24]. We hypothesize that the DDAG neurons function as a conduit between the descending neurons, vpoDN and DNp13, and the motor circuits for vaginal plate opening and ovipositor extrusion, respectively [23,24]. We have shown that optogenetic activation of the DDAG neurons triggers vaginal plate opening or ovipositor extrusion depending upon the female’s mating status. How the activity of the DDAG neurons is integrated with female mating state is currently unclear. A recent study showed that *ppk+* mechanosensory neurons in the female reproductive tract sense ovulation and their activity permits the DNp13 neurons to engage the motor circuits for ovipositor extrusion [24]. One possibility is that vaginal plate opening and ovipositor extrusion are controlled by distinct subtypes of DDAG neurons. The DDAG subtypes that influence ovipositor extrusion may function downstream of the DNp13 neurons but upstream of the site at which *ppk+* sensory neurons integrate with the neural circuit for ovipositor extrusion.

*Dsf* encodes an orphan nuclear receptor homologous to the *Drosophila tailless* (*tll*) and vertebrate *Tlx* genes [13]. In mice, *Tlx* is expressed in all adult neural stem cells [33], where it functions to maintain the cells in a proliferative state [34,35]. Similarly, in *Drosophila, tll* regulates the maintenance and proliferation of most neural progenitors in the protocerebrum [36]. At the developmental stages we examined, *dsf*^Ga14^ expression in the ventral nerve cord was observed exclusively in mature neurons and not in any neuronal progenitors. This suggest that *dsf* may regulate neuron number post-mitotically, directing the survival or apoptosis of female-specific DDAG neurons depending upon expression of *dsx*M.

How might DSX-M and DSF interact molecularly? Although nuclear receptors including *dsf* generally act as transcriptional repressors, some can also function as activators upon binding to a ligand [37–39] or after post-translational modification [40]. Thus, in one scenario, the expression of DSX-M could sex-specifically impact DSF activity by regulating the generation of DSF’s ligand (if it has one) or the occurrence of a post-translational modification. Alternatively (but not exclusively), expression of *dsf* may be directly regulated by DSX proteins. Consistent with this scenario, an *in vivo* genome-wide study of DSX occupancy identified *dsf* as a putative DSX target gene [41]. It is therefore possible that DSX-M sex-specifically regulates *dsf* expression in a subset of DDAG neurons that contribute to female-specific abdominal behaviors, thereby directing the removal of the neurons in males but not females.

## Experimental Procedures

### Fly stocks

Fly stocks were maintained on standard cornmeal and molasses food at 25°C. The stocks used in this study include the following: Canton S, *w^1118^, Df(2L)ED284/CyO* (BDSC 8040), *Df(2L)BSC183/CyO* (BDSC 9611), *pJFRC32-10XUAS-IVS-nlsGFP* (attP40), *pJFRC29-10XUAS-IVS-myr::GFP-p10* (attP2), *dsx*^LexA::p65^ [20], *pJFRC79-8XLexAop2-FlpL* (attP40), *pJFRC41-10XUAS-FRT>STOP>FRT-myr::gfp* (su(Hw)attP1), *20XUAS-FRT>STOP>FRT-CsChrimson-mVenus* (VK5), *UAS-FRT>STOP>FRT-tnt.e* and *UAS-FRT>STOP>FRT-tnt.QA, dsx*^Ga15(Δ2)^ [32], *P{y[+t7.7] v[+t1.8]=TRiP.GLV21010}attP2 (i.e., UAS-dsx_ShmiR;* obtained from *Drosophila* Transgenic RNAi Project at Harvard Medical School)*, UAS-traF* (BDSC 4590), and *UAS-P35^BH1^* (BDSC 5072). Sex Peptide null (SP^0^) males were generated by crossing *Δ130/TM3* males to *0325/TM3* females (gift from M. Wolfner, Cornell). *UAS-dsf_ShmiR* (attP2) was created according to protocols described in Haley *et al.* [42]. Two stem-loops targeting sequences in *dsf*’s fourth exon were placed into an intron from the *ftz* gene upstream of a cDNA encoding *nls::gfp.* This construct was cloned into the BglII/XbaI sites in pMUH.

### Construction of *dsx*^Ga14^ and *dsf*^Del^ alleles

The *dsf*^Ga14^ allele was created using CRISPR/Cas9-mediated homology-directed repair to precisely introduce Gal4 into *dsfs* start codon in exon 1. The Gal4 cDNA/hsp70 yeast terminator sequence was amplified from pBPGuW (Addgene). A donor plasmid was generated by combining a 1,050-bp left homology arm, the Gal4/terminator sequence, 3XP3::DsRed marker (with an inverted orientation), a 1,149-bp right homology arm, and a 1.8-kb backbone using Gibson Assembly [43]. Guide RNAs (5’-CATCAACGGAAAATGGGCAC-3’ and 5’-ATACTTCCATCAACGGAAAA-3’) were cloned into the pCFD4 plasmid [44]. The donor and pCFD4 plasmids, *in vitro* transcribed codon optimized Cas9 mRNA, and a lig4 siRNA were coinjected into *w*^1118^ embryos. The *dsf*^Del^ allele was created using an identical strategy except the Gal4/terminator sequence was excluded from the donor plasmid.

### *in situ* hybridization chain reaction and immunohistochemistry

Nervous systems were dissected in 1X PBS, fixed in 4% paraformaldehyde buffered in PBS for 35 min at room temperature, and rinsed and washed in PBT (PBS with 1% Triton X-100). *in situ* hybridization chain reaction experiments were conducted as described in [17] using HCR probe-sets *(dsf-B1,* Lot #: PRF061) and hairpins (B1h1-AF488, B1h2-AF488) synthesized by Molecular Instruments™. We used a probe-set size of 20. After dissection and fixation, the tissues were pre-hybridized in prewarmed probe hybridization buffer (Molecular Instruments™) for 30 minutes at 37°C and incubated with HCR probes in probe hybridization buffer overnight at 37°C. Tissues were washed the next day in prewarmed probe wash buffer four times, 15 minutes each at 37°C, and washed in 5X SSCT (UltraPure 20X SSC Buffer, Invitrogen #15557044, diluted in water) three times, 5 minutes at room temperature. Tissues were pre-amplified in amplification buffer (Molecular Instruments™) for 30 minutes at room temperature and incubated with snap-cooled HCR hairpins in amplification buffer overnight at room temperature, protected from light. Tissues were then washed with 5X SSCT at room temperature twice for 5 minutes, twice for 30 minutes, and once for 5 minutes before being mounted in Vectashield (Vector Laboratories, H-100) or proceeding with an immunohistochemistry protocol. Immunohistochemistry was performed as described in [29]. Nervous systems were incubated with primary antibodies in PBT overnight at 4°C. Tissues were washed in PBT at room temperature for several hours and incubated overnight with secondary antibodies at 4°C. Tissues were washed in PBT for several hours at room temperature, mounted onto poly-lysine coated coverslips, dehydrated through an ethanol series, cleared in xylenes, and mounted in DPX (Sigma-Aldrich) on a slide. If immunohistochemistry followed *in situ* HCR, nervous systems were mounted in Vectashield. The following antibodies were used: rabbit anti-GFP (Invitrogen #A11122; 1:1,000), rat anti-DN-cadherin (DN-Ex #8, Developmental Studies Hybridoma Bank; 1:50), mouse anti-Elav (9F8A9, Developmental Studies Hybridoma Bank; 1:20), AF-647 goat anti-rat (Invitrogen #A21247; 1:500), Fluorescein (FITC)-conjugated donkey anti-rabbit (Jackson ImmunoResearch; 1:500), AF-568 goat anti-mouse (Invitrogen #A11004; 1:500). Tissues were imaged on a Leica SP8 confocal microscope at 40X with optical sections at 0.3 μm intervals.

### Behavior and optogenetic assays

Newly eclosed flies were collected under CO2 anesthesia and aged for 7–10 days at 25°C and ~50% humidity in a 12-hr light/dark cycle unless otherwise noted. Virgin females and males used in optogenetic assays were raised in darkness at 25°C on food containing 0.2 mM *all-trans-* Retinal (Sigma-Aldrich) before and after eclosion, grouped in vials according to sex, and aged for 10–14 days. Flies were chilled on ice and decapitated under a low intensity of blue light (470 nm) and allowed to recover for ~30 min before being transferred into behavioral chambers (diameter: 10 mm, height: 3 mm). The chambers were kept in darkness and placed above an LED panel that provided infrared light (850 nm) for continuous illumination and photoactivation with red light (635 nm). The intensity and pattern of red-light illumination were controlled by a customized Arduino script. Light intensity was confirmed with an optical power meter (Thorlabs, PM160). A camera (FLIR Blackfly S USB3, BFS-U3-31S4M-C) equipped with a long-pass filter (cut-on wavelength of 800 nm; Thorlabs, FEL0800) was placed above the chamber and videos were recorded at 55 fps with a resolution of 0.03 mm/pixel. Courtship and receptivity assays were conducted using group- and singly-housed virgin females and males, respectively. Behavior assays were done at 25°C under white light, ~1-2 hrs after the start of the subjective day. Individual females and naive males were loaded into behavioral chambers (diameter: 10 mm, height: 3 mm) and recorded for 30 min using a Sony Vixia HFR700 video camera. Ovipositor extrusion was assessed using mated females prepared 24 hrs before the assay by grouping 10 virgin females and 20 wild-type Canton S or *SP*^0^ in a vial. Mated females were anesthetized on ice and sorted 30 min before the assay. Individual females and naive Canton S males were loaded into behavioral chambers as described above and video recorded for 15 min. For male abdominal curling measurements, virgin Canton S females and experimental or control males were collected and housed as described above. Virgin females were anesthetized on ice and decapitated to prevent copulation 30 min before being transferred individually into behavioral chambers with a male. The pairs were recorded for 15 min. Change in abdominal length was measured in ImageJ using frames before and during vaginal plate opening or ovipositor extrusion in which the female was in the same position in the chamber. A ruler was included in the video and used to set the scale of measurement in mm. The length of abdomen was measured by drawing a line from the base of scutellum to the posterior tip of the female’s abdomen before and during vaginal plate opening or ovipositor extrusion. The change in abdominal length was determined by calculating the difference between the two measurements. The vaginal plate opening and abdominal curling indices were calculated by dividing the amount of time the female opened her vaginal plates or the male curled his abdomen by the total photoactivation time of each bout. Courtship index was measured as the total amount of time the male spent performing any courtship behavior divided by the total observation time. All p-values were measured in Matlab.

## Acknowledgements

We thank Y. Ding, J. Lillvis, and E. Preger-Ben Noon for helpful discussions, and A. McStravog for administrative assistance. DLS is an Investigator with Howard Hughes Medical Institute. TRS is supported by a grant from the National Science Foundation.

---

**Supplemental Figure 1.** *dsf*^Ga14^ expression in the CNS of larval, pupal, and adult flies. (A-D) *dsf*^Ga14^-labeled cells in the CNS of females and males express the neuronal marker, ELAV. (E–F) *dsf*^Ga14^ labels neurons in the late third instar larval female and male CNS. (G–J) *dsf*^Ga14^ labels CNS neurons in females and males during pupal life. (K) In the abdominal ganglion of *dsf*^Ga14^ > *UAS-myr::gfp* females and males, *dsf* hybridization signals were found exclusively in ELAV+ neurons that are GFP+ indicating that *dsf*^Ga14^ accurately targets all *dsf*-expressing neurons. *dsf*^Ga14^ activity was not observed in any motoneurons or sensory neurons.

**Supplemental Figure 2.** NMJs of *dsf* mutant females and females expressing tetanus toxin in the DDAG neurons. (A-E) Uteri from females of the corresponding genotypes were stained with an anti-HRP antibody to visualize the synapses at the uterine wall. *Dsf* null females lack uterine synapses, as reported in ref. [12], as do and *dsf*^Ga14^/*dsf*^DEL^ females. Females expressing *tnt.e* in the DDAG neurons do not exhibit any obvious defects in synaptic morphology relative to *tnt.QA-* expressing control females.

**Supplemental Figure 3.** Knock-down of *dsx* and *dsf* transcripts in the DDAG neurons of females and males. (A-C) Mating rate (A), frequency of vaginal plate opening (vpo) behavior (B), and number of eggs laid in 20-22 hrs (C) of *dsf*^Ga14^ > *UAS-dsx_ShmiR* and control females. (D, E) Mating rate (D) and frequency of abdominal curls (E) of *dsf*^Ga14^ > *UAS-dsx_ShmiR* and control males. (F-H) Mating rate (F), frequency of vpo (G), and number of eggs laid in 20-22 hrs (H) of *dsx*^Ga14^ > *UAS-dsf_ShmiR* and control females. (I, J) Mating rate (I) and frequency of abdominal curls (J) of *dsx*^Ga14^ > *UAS-dsf_ShmiR* and control males. Significance (P<0.05) in (A, D, F, I) was measured with a Logrank test. (B, C, E, G, H, J) show individual points, mean, and SD. Significance was measured with a Rank Sum test. Same letter denotes no significant difference.

**Supplemental Video 1.** Optogenetic activation of DDAG neurons in *dsf*^Ga14^/+ virgin females. Photoactivation period begins with the green dot.

**Supplemental Video 2.** Optogenetic activation of DDAG neurons in *dsf*^Ga14^/+ females that mated with Canton S males.

**Supplemental Video 3.** Optogenetic activation of DDAG neurons in *dsf*^Ga14^/+ females that mated with Sex Peptide null males.

**Supplemental Video 4.** Optogenetic activation of DDAG neurons in *dsf*^Ga14^/+ males. **Supplemental Video 5.** Optogenetic activation of DDAG neurons in *dsf*^Ga14^ > *UAS-dsx-ShmiR* virgin females.

**Supplemental Video 6.** Optogenetic activation of DDAG neurons in *dsf*^Ga14^ > *UAS-dsx-ShmiR* males.

**Supplemental Video 7.** Optogenetic activation of DDAG neurons in *dsf*^Ga14^/*dsf*^Del^ virgin females.

**Supplemental Video 8.** Optogenetic activation of DDAG neurons in *dsf^Ga14^/dsf^Del^* males.

## References

1. Billeter, J.-C., Rideout, E.J., Dornan, A.J., and Goodwin, S.F. (2006). Control of male sexual behavior in Drosophila by the sex determination pathway. Curr. Biol. 16, R766–R776.

2. Manoli, D.S., Foss, M., Villella, A., Taylor, B.J., Hall, J.C., and Baker, B.S. (2005). Male-specific fruitless specifies the neural substrates of Drosophila courtship behaviour. Nature 436, 395–400.

3. Demir, E., and Dickson, B.J. (2005). fruitless splicing specifies male courtship behavior in Drosophila. Cell 121, 785–94.

4. Stockinger, P., Kvitsiani, D., Rotkopf, S., Tirin, L., and Dickson, B.J. (2005). Neural circuitry that governs Drosophila male courtship behavior. Cell 121, 795–807.

5. Robinett, C.C., Vaughan, A.G., Knapp, J.M., and Baker, B.S. (2010). Sex and the single cell. II. there is a time and place for sex. PLoS Biol. 8(5):e1000365.

6. Rideout, E.J., Dornan, A.J., Neville, M.C., Eadie, S., and Goodwin, S.F. (2010). Control of sexual differentiation and behavior by the doublesex gene in Drosophila melanogaster. Nat. Neurosci. 13, 458–66.

7. Waterbury, J.A., Jackson, L.L., and Schedl, P. (1999). Analysis of the doublesex female protein in Drosophila melanogaster: Role in sexual differentiation and behavior and dependence on intersex. Genetics. 152, 1653–67.

8. Shirangi, T.R., Taylor, B.J., and McKeown, M. (2006). A double-switch system regulates male courtship behavior in male and female Drosophila melanogaster. Nat Genet. 38, 1435–;1439.

9. Kvitsiani, D., and Dickson, B.J. (2006). Shared neural circuitry for female and male sexual behaviours in Drosophila. Curr. Biol. 16(10):R355–6.

10. Shirangi, T.R., and McKeown, M. (2007). Sex in flies: What “body-mind” dichotomy? Dev. Biol. 306, 10–19.

11. Villella, A., and Hall, J.C. (2008). Chapter 3 Neurogenetics of Courtship and Mating in Drosophila. Adv. Genet. 62, 167–184.

12. Finley, K.D., Taylor, B.J., Milstein, M., and McKeown, M. (1997). *dissatisfaction*, a gene involved in sex-specific behavior and neural development of *Drosophila melanogaster*. PNAS 94, 913–8.

13. Finley, K.D., Edeen, P.T., Foss, M., Gross, E., Ghbeish, N., Palmer, R.H., Taylor, B.J., and McKeown, M. (1998). dissatisfaction encodes a tailless-like nuclear receptor expressed in a subset of CNS neurons controlling Drosophila sexual behavior. Neuron 21, 1363–1374.

14. Choi, H.M.T., Schwarzkopf, M., Fornace, M.E., Acharya, A., Artavanis, G., Stegmaier, J., Cunha, A., and Pierce, N.A. (2018). Third-generation in situ hybridization chain reaction: multiplexed, quantitative, sensitive, versatile, robust. Development 145(12):dev165753.

15. Choi, H.M.T., Beck, V.A., and Pierce, N.A. (2014). Next-Generation in Situ Hybridization Chain Reaction: Higher Gain, Lower Cost, Greater Durability. Zebrafish. 11, 488–9.

16. Choi, H.M.T., Chang, J.Y., Trinh, L.A., Padilla, J.E., Fraser, S.E., and Pierce, N.A. (2010). Programmable in situ amplification for multiplexed imaging of mRNA expression. Nat. Biotechnol. 28, 1208–1212.

17. Duckhorn, J.C., Junker, I., Ding, Y., and Shirangi, T.R. (2021). Combined in situ hybridization chain reaction and immunostaining to visualize gene expression in whole-mount Drosophila central nervous systems. bioRxiv. doi: doi.org/10.1101/2021.08.09.455671

18. Court, R., Namiki, S., Armstrong, J.D., Börner, J., Card, G., Costa, M., Dickinson, M., Duch, C., Korff, W., Mann, R., et al. (2020). A Systematic Nomenclature for the Drosophila Ventral Nerve Cord. Neuron 107, 1071–1079.e2.

19. Ito, K., Shinomiya, K., Ito, M., Armstrong, J.D., Boyan, G., Hartenstein, V., Harzsch, S., Heisenberg, M., Homberg, U., Jenett, A., et al. (2014). A systematic nomenclature for the insect brain. Neuron 81, 755–765.

20. Zhou, C., Pan, Y., Robinett, C.C., Meissner, G.W., and Baker, B.S. (2014). Central brain neurons expressing doublesex regulate female receptivity in Drosophila. Neuron 83, 149–163.

21. Klapoetke, N.C., Murata, Y., Kim, S.S., Pulver, S.R., Birdsey-Benson, A., Cho, Y.K., Morimoto, T.K., Chuong, A.S., Carpenter, E.J., Tian, Z., et al. (2014). Independent optical excitation of distinct neural populations. Nat. Methods 11, 338–46.

22. Mezzera, C., Brotas, M., Gaspar, M., Pavlou, H.J., Goodwin, S.F., and Vasconcelos, M.L. (2020). Ovipositor Extrusion Promotes the Transition from Courtship to Copulation and Signals Female Acceptance in Drosophila melanogaster. Curr. Biol. 30, 3736–3748.e5.

23. Wang, K., Wang, F., Forknall, N., Yang, T., Patrick, C., Parekh, R., and Dickson, B.J. (2021). Neural circuit mechanisms of sexual receptivity in Drosophila females. Nature 589, 577–581.

24. Wang, F., Wang, K., Forknall, N., Parekh, R., and Dickson, B.J. (2020). Circuit and Behavioral Mechanisms of Sexual Rejection by Drosophila Females. Curr. Biol. 30, 3749–3760.e3.

25. Kimura, K.I., Sato, C., Koganezawa, M., and Yamamoto, D. (2015). Drosophila ovipositor extension in mating behavior and Egg deposition involves distinct sets of brain interneurons. PLoS One 10(5):e0126445.

26. Kubli, E. (1992). The sex-peptide. Bioessays 14, 779–784.

27. Sweeney, S.T., Broadie, K., Keane, J., Niemann, H., and O’Kane, C.J. (1995). Targeted expression of tetanus toxin light chain in Drosophila specifically eliminates synaptic transmission and causes behavioral defects. Neuron 14, 341–351.

28. Haley, B., Hendrix, D., Trang, V., and Levine, M. (2008). A simplified miRNA-based gene silencing method for Drosophila melanogaster. Dev. Biol. 321, 482–90.

29. Shirangi, T.R., Wong, A.M., Truman, J.W., and Stern, D.L. (2016). Doublesex Regulates the Connectivity of a Neural Circuit Controlling Drosophila Male Courtship Song. Dev. Cell 37, 533–544.

30. Hay, B.A., Wolff, T., and Rubin, G.M. (1994). Expression of baculovirus P35 prevents cell death in Drosophila. Development 120, 2121–2129.

31. McKeown, M., Belote, J.M., and Boggs, R.T. (1988). Ectopic expression of the female transformer gene product leads to female differentiation of chromosomally male drosophila. Cell 53, 887–895.

32. Pan, Y.F., Robinett, C.C., and Baker, B.S. (2011). Turning Males On: Activation of Male Courtship Behavior in Drosophila melanogaster. PLoS One 6(6):e21144.

33. Shi, Y., Lie, D.C., Taupin, P., Nakashima, K., Ray, J., Yu, R.T., Gage, F.H., and Evans, R.M. (2004). Expression and function of orphan nuclear receptor TLX in adult neural stem cells. Nature 427, 78–83.

34. Liu, H.K., Belz, T., Bock, D., Takacs, A., Wu, H., Lichter, P., Chai, M., and Schütz, G. (2008). The nuclear receptor tailless is required for neurogenesis in the adult subventricular zone. Genes Dev. 22, 2473–2478.

35. Zhang, C.L., Zou, Y., He, W., Gage, F.H., and Evans, R.M. (2008). A role for adult TLX-positive neural stem cells in learning and behaviour. Nature 451, 1004–1007.

36. Younossi-Hartenstein, A., Green, P., Liaw, G.J., Rudolph, K., Lengyel, J., and Hartenstein, V. (1997). Control of early neurogenesis of the Drosophila brain by the head gap genes tll, otd, ems, and btd. Dev. Biol. 182, 270–283.

37. Mangelsdorf, D.J., and Evans, R.M. (1995). The RXR heterodimers and orphan receptors. Cell 83, 841–850.

38. Pitman, J.L., Tsai, C.C., Edeen, P.T., Finley, K.D., Evans, R.M., and McKeown, M. (2002). DSF nuclear receptor acts as a repressor in culture and in vivo. Dev. Biol. 245, 315–28.

39. Lee, J.W., Lee, Y.C., Na, S.Y., Jung, D.J., and Lee, S.K. (2001). Transcriptional coregulators of the nuclear receptor superfamily: Coactivators and corepressors. Cell. Mol. Life Sci. 58, 289–297.

40. Hammer, G.D., Krylova, I., Zhang, Y., Darimont, B.D., Simpson, K., Weigel, N.L., and Ingraham, H.A. (1999). Phosphorylation of the nuclear receptor SF-1 modulates cofactor recruitment: Integration of hormone signaling in reproduction and stress. Mol. Cell 3, 521–526.

41. Clough, E., Jimenez, E., Kim, Y.A., Whitworth, C., Neville, M.C., Hempel, L.U., Pavlou, H.J., Chen, Z.X., Sturgill, D., Dale, R.K., et al. (2014). Sex- and tissue-specific functions of drosophila doublesex transcription factor target genes. Dev. Cell 31, 761–773.

42. Haley, B., Foys, B., and Levine, M. (2010). Vectors and parameters that enhance the efficacy of RNAi-mediated gene disruption in transgenic Drosophila. PNAS 107, 11435–11440.

43. Gibson, D.G., Young, L., Chuang, R.-Y., Venter, J.C., Hutchison, C.A., and Smith, H.O. (2009). Enzymatic assembly of DNA molecules up to several hundred kilobases. Nat. Methods 6, 343–5.

44. Port, F., Chen, H., Lee, T., and Bullock, S.L. (2014). Optimized CRISPR / Cas tools for efficient germline and somatic genome engineering in Drosophila. PNAS 111, E2967–76.

