## Supplemental Data 1 for "Regulation of sexually dimorphic abdominal courtship behaviors in *Drosophila* by the *Tlx/tailless*-like nuclear receptor, *Dissatisfaction*"

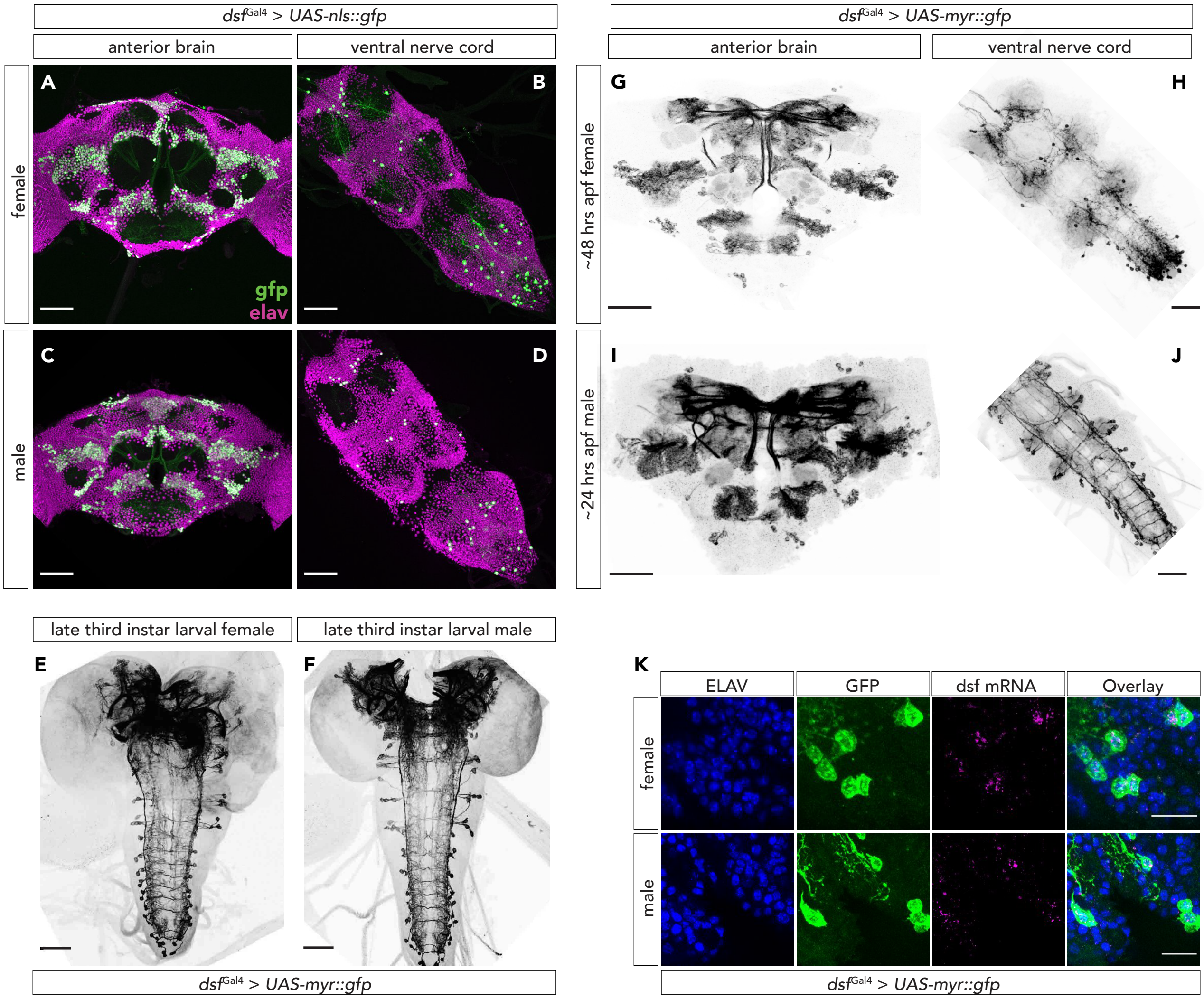


**Supplemental Figure 1.** dsfGal4 expression in the CNS of larval, pupal, and adult flies.
(A–D) DsfGal4-labeled cells in the CNS of females and males express the neuronal marker, ELAV.
(E, F) dsfGal4 labels neurons in the late third instar larval female and male CNS.
(G–J) dsfGal4 labels CNS neurons in females and males during pupal life.
(K) In the abdominal ganglion of dsfGal4 > UAS-myr::gfp females and males, dsf hybridization signals were found exclusively in ELAV+ neurons that are GFP+ indicating that dsfGal4 accurately targets all dsf-expressing neurons. dsfGal4 activity was not observed in any motoneurons or sensory neurons.

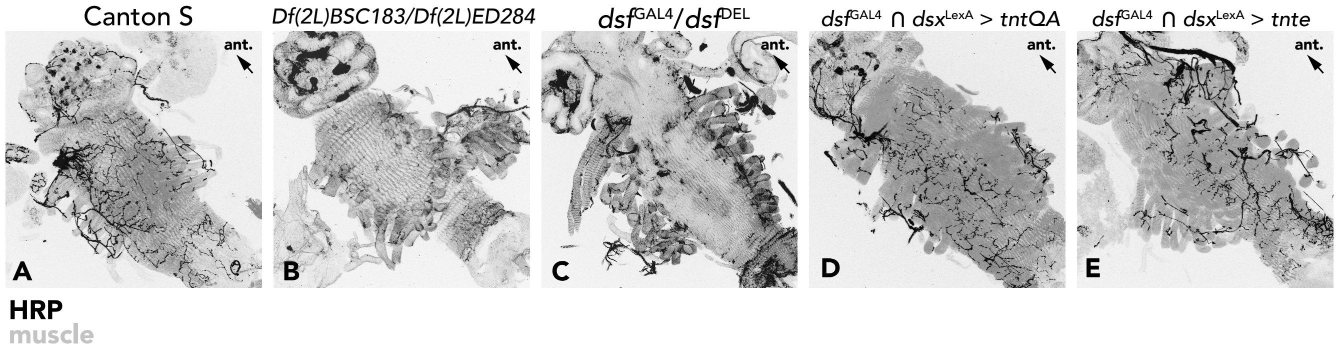


**Supplemental Figure 2.** NMJs of dsf mutant females and females expressing tetanus toxin in the DDAG neurons. Uteri from females of the corresponding genotypes were stained with an anti-HRP antibody to visualize the synapses at the uterine wall. Dsf null females lack uterine synapses, as reported in ref. [12], as do and dsfGal4/dsfDEL females. Females expressing tnt.e in the DDAG neurons do not exhibit any obvious defects in synaptic morphology relative to tnt.QA-expressing control females.

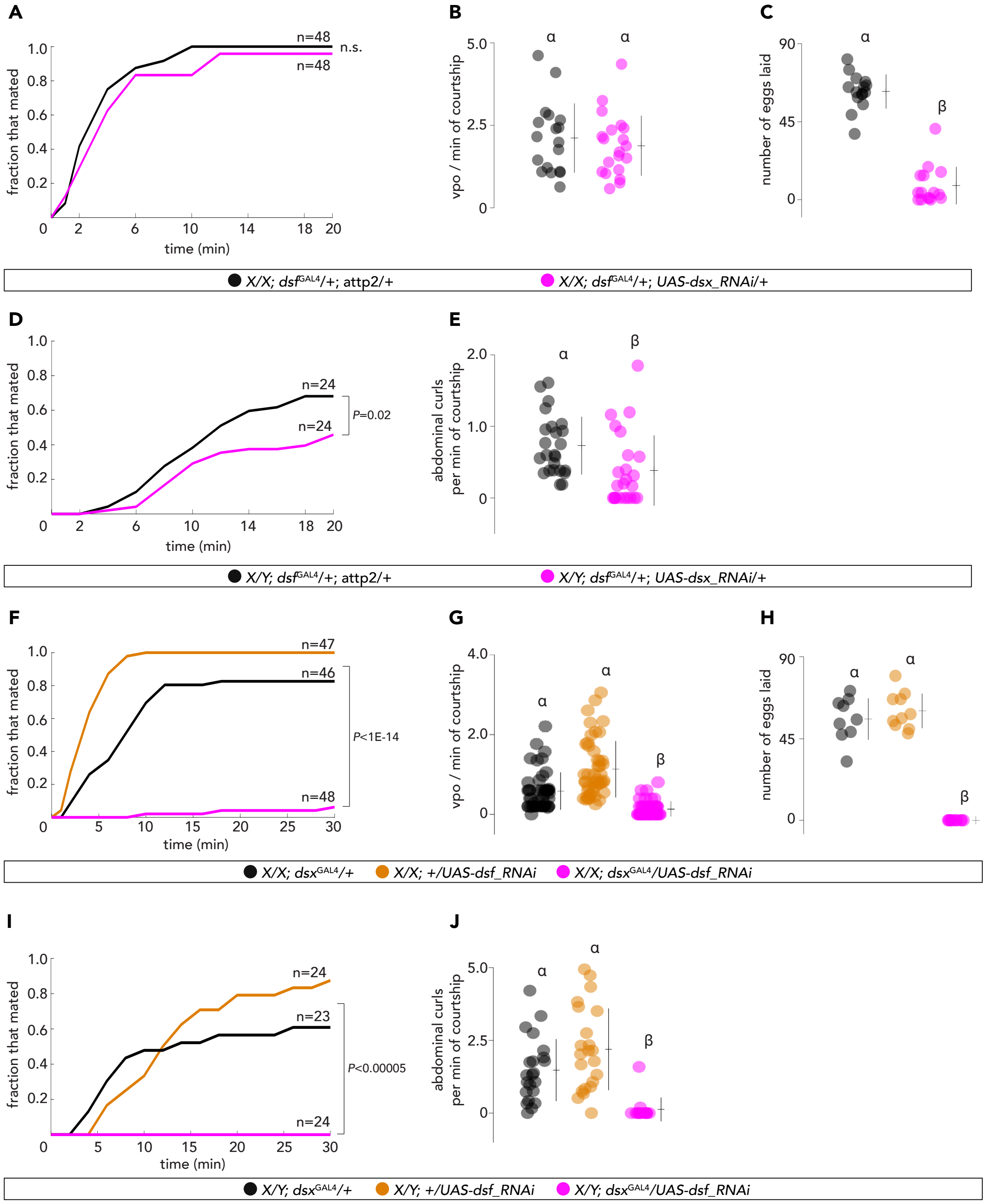


**Supplemental Figure 3.** Knock-down of dsx and dsf transcripts in the DDAG neurons of females and males.
(A–C) Mating rate (A), frequency of vaginal plate opening (vpo) behavior (B), and number of eggs laid in 20–22 hrs (C) of dsfGal4 > UAS-dsx_ShmiR and control females.
(D, E) Mating rate (D) and frequency of abdominal curls (E) of dsfGal4 > UAS-dsx_ShmiR and control males.
(F–H) Mating rate (F), frequency of vpo (G), and number of eggs laid in 20–22 hrs (H) of dsxGal4 > UAS-dsf_ShmiR and control females.
(I, J) Mating rate (I) and frequency of abdominal curls (J) of dsxGal4 > UAS-dsf_ShmiR and control males.
Significance ($P<0.05$) in (A, D, F, I) was measured with a Logrank test. (B, C, E, G, H, J) show individual points, mean, and SD. Significance was measured with a Rank Sum test. Same letter denotes no significant difference.
